# Monkeys have rhythm

**DOI:** 10.1101/2024.03.11.584468

**Authors:** Vani G. Rajendran, Juan Pablo Marquez, Luis Prado, Hugo Merchant

## Abstract

Synchronizing movements to music is one of the hallmarks of human culture whose evolutionary and neurobiological origins remain unknown. The ability to synchronize movements requires 1) detecting a steady rhythmic pulse, or beat, out of a stream of complex sounds, 2) projecting this rhythmic pattern forward in time to predict future input, and 3) timing motor commands in anticipation of predicted future beats. Here, we demonstrate that the macaque is capable of synchronizing taps to a subjective beat in real music, and even spontaneously chooses to do so over alternative strategies. This contradicts the influential “vocal learning hypothesis” that musical beat synchronization is only possible in species with complex vocalizations such as humans and some songbirds. We propose an alternative view of musical beat perception and synchronization ability as a continuum onto which a wider range of species can be mapped depending on their ability to perform and coordinate the general abilities listed above through association with reward.

## Introduction

Creating and appreciating music is an amazing human capability. Despite substantial differences in harmonic and rhythmic preferences between human cultures [1, 2], music is a nearly universal cultural phenomenon [3–5] whose evolutionary origin remains mysterious [6, 7] and highly debated [8, 9].

Neuroscientists have approached this question through comparative studies across species to explore the repertoire of natural and trained behaviors that can be evoked by human music and their biological implications [10]. A key behavior exhibited by humans is the spontaneous perception of a steady pulse, or beat, in music – and the synchronization of movements to these beats [11–14]. This ability requires detecting an abstract rhythmic pattern out of a continuous stream of complex sound, projecting this pattern forward into the future to predict when the next beat is likely to occur [15], and timing the initiation of motor commands such that movements coincide with, and in fact usually anticipate [16], the next beat. Auditory-motor synchronization to beat has been observed from preschool age in humans [17], with rhythmic engagement with music from infancy itself [18, 19]. Interestingly, auditory-motor synchronization has only been observed in some bird species [20, 21] and occasional exceptional individuals of other species [22], leaving a gap in our understanding of both the evolutionary and neurobiological basis of the capacity for rhythmic synchronization to music.

One influential hypothesis in the field is the vocal learning and rhythmic synchronization hypothesis [23, 24], which states that high vocal learning is a preadaptation for the spontaneous perception and synchronization to musical beat. This hypothesis is supported by the observation that the most musical species do tend to also have complex vocal communication, i.e. humans and songbirds. The strongest challenge to date of the vocal learning hypothesis is a sea lion who shows a flexible ability to synchronize to music [22]. However, it has been argued that even though the sea lion is not traditionally considered a vocal-learning species, it still shows a high degree of vocal flexibility [24]. Further comparisons show that even synchronization to explicit beats such as in the case of a metronome are better performed by vocal-learning species [25]. So though existing evidence tends to favor this hypothesis, this perspective has left open the question of how human musicality could have evolved along the primate line, and has limited lab-based studies of musical beat perception to largely bird species, whose neurophysiology and skeletomotor systems differ vastly from those of humans.

The macaque, a non vocal-learning primate, is a widely used animal model that has the advantage of having both neuroanatomical and skeletomotor function similar to humans. It was thought that macaques do not share human beat synchronization ability due to their taps naturally lagging metronome beats when attempting to synchronize [26, 27]. However, macaques are able to produce human-like predictive and tempo-flexible synchronization to metronome beats when reward was made contingent on the asynchrony between taps and beats [28], indicating that predictive synchronization can be trained in a non vocal-learning species. As a result of establishing the macaque as a model of predictive auditory-motor synchronization to explicit metronome beats, extracellular recordings performed in awake, behaving macaques are beginning to reveal in unprecedented detail the neural activity patterns and dynamics that underlie this ability [29–32]. Crucially, these findings enable direct comparison with human behavior and neurophysiology during perception and synchronization to explicit metronome beats [27], and thereby begin bridging the neurophysiological and evolutionary void in our understanding of beat perception.

Here, we push this one important step further by demonstrating that macaques can also perceive and tap to a *subjective* beat in real music, and even choose to do so when they can ignore the sound or produce intervals of their own choosing for reward. Comparison with human tapping reveals fundamental similarities and differences that point to key neurobiological similarities and differences that may produce this pattern of results. In this context, we propose that musical beat perception and synchronization may be a spectrum onto which a wider range of species may be mapped based on the extent of their ability to perform and coordinate four key component processes of this ability (auditory pattern detection, prediction, auditory-motor feedback, and reward-based reinforcement learning – the “Four Components (4Cs)” hypothesis).

## Results

### 1. Macaques tap a subjective beat that is linked to features in real music

Two adult male macaques, previously trained to synchronize to explicit metronome beats, were trained to abstract from explicit beats to a subjective beat in music (see Methods). The subjects initiated a trial by placing their hand on a holding bar (Fig. 1a-b), upon which a yellow square would appear on the screen and the music would start. The disappearance of the yellow square would signal to the monkey to begin tapping (Fig. 1c). In *Experiment 1: Music*, three different pieces of music were presented (see Methods). Spectrograms of these musical excerpts are shown in Fig. 1e. The target intervals that the monkeys needed to tap in order to receive reward corresponded to the tempi of the songs used as reported in [33] (465 ms, 732 ms, and 882 ms, corresponding to tempi of 2.2, 1.4, and 1.1 Hz, or 129, 82, and 68 beats per minute, respectively; see Supplementary Audio A1-A3). Songs (and therefore, tempi) were pseudo-randomly presented from trial to trial.

The subjective interpretation of an isochronous beat expressed through tapping has both a period, marked by the inter-tap intervals (ITIs), and a phase. Crucially, to receive reward, the monkey only needed to produce five taps where the ITIs were within +/-20% of the target interval (see Methods). In other words, the monkeys were never explicitly trained or rewarded to produce any particular phase. Despite this, both monkeys produced a consistent tapping phase for all three musical excerpts (p<0.001, Rayleigh test for uniformity, N=138-320 trials per subject; see Supplementary Videos V1-V2). To test whether the monkeys were truly synchronizing to some feature in the music (rather than simply producing a stereotyped response triggered by the visual go cue), the go signal was shifted by pi with respect to the features in the music (Fig. 1d), while maintaining the same trial duration. If the monkeys were simply producing a stereotyped response triggered by the visual go cue, the phase of tapping with respect to the go cues should align between the original and pi-shifted music. Figure 1f illustrates that tapping phase distributions are significantly different between original (“0”) and pi-shifted (“π”) songs (p<0.05, Watson’s U^2^ test, N=270-586 trials, Bonferroni-corrected for multiple comparisons), with the exception of M2 for the medium tempo song, suggesting strongly that monkeys are indeed synchronizing to some feature in the stimulus. This difference in tapping phase emerges for both monkeys by the second or third tap in the sequence (Supplementary Figs. S1-S2) and is visually clearer overall if trials with ITIs exceeding +/-15% are excluded (Supplementary Fig. S3). Even in the exceptional case, M2 appears to be synchronizing to a feature that is in antiphase with his interpretation for the original music (see Supplementary Videos S3), which also aligns with some human interpretations of beat for this excerpt (see Supplementary Fig. S4). Overall, monkeys show a less consistent shift in their tapping than the roughly consistent shift by pi observed in humans.

**Fig. 1.**
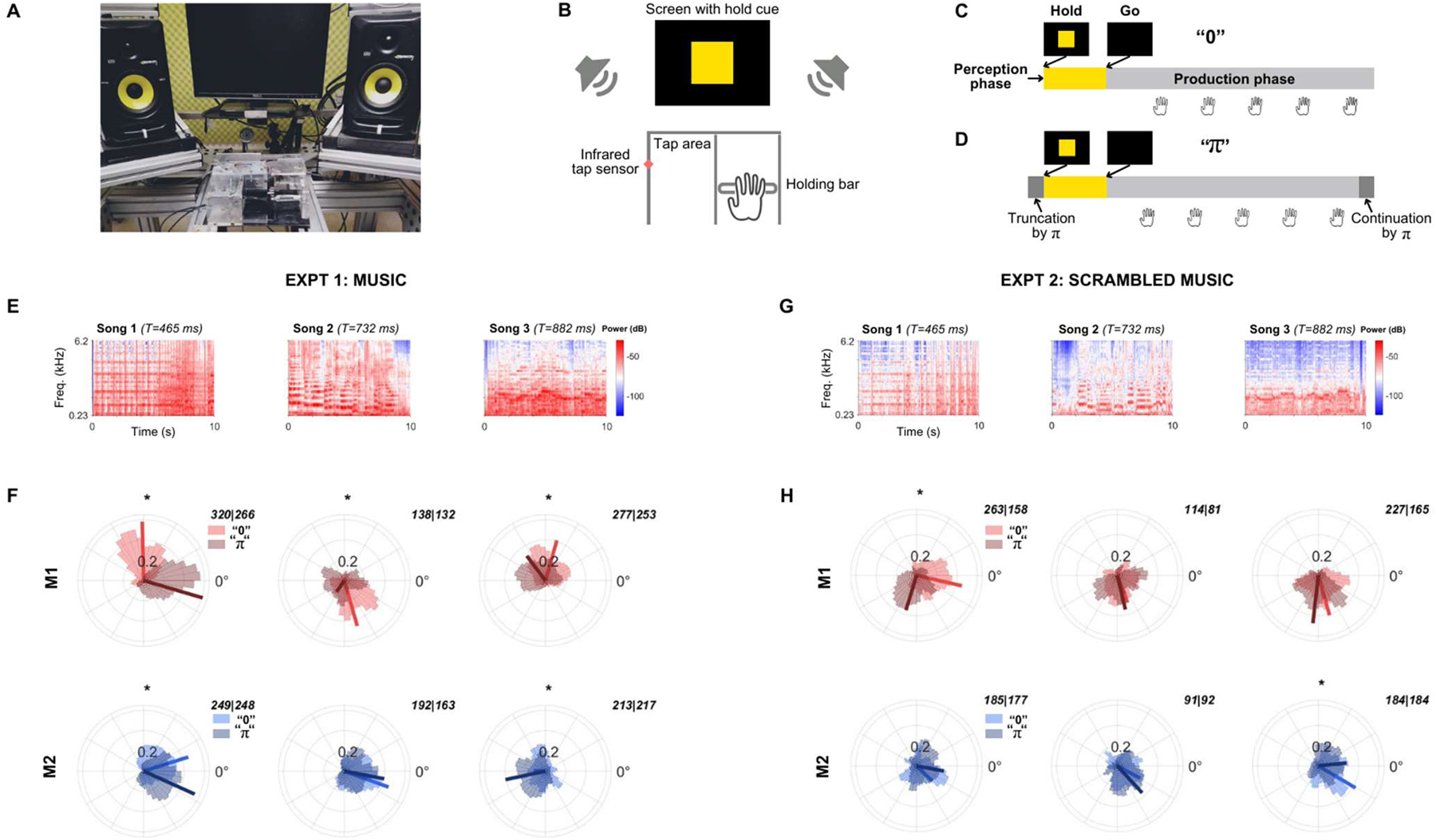
Macaques synchronize their taps to music. **a)** Photo of experimental setup from the experimental subject’s perspective. **b)** Schematic illustration showing details of the setup. Note that going from the holding bar to the tap area to produce the first tap requires a relatively substantial movement compared to subsequent taps performed inside the tap area. **c)** Schematic illustration of a trial, which begins with a perception phase, after which subjects needed to produce five correct taps to receive reward. **d)** Schematic illustration of a pi-shifted control, which involves truncating the start of the stimulus such that the Hold and Go cues are shifted with respect to features in the stimulus. **e) S**pectrograms of the musical excerpts used in Experiment 1. **f)** Tapping responses for *Experiment 1: Music* for M1 (top row) and M2 (bottom row), with Songs 1-3 arranged from left to right. Each circular histogram shows tapping responses to the original “0” (in color) and pi-shifted “π” (same color but grayed) conditions. 0° corresponds to the consensus tapping phase of human participants in this study for each of the original 3 musical excerpts (see Supplementary Fig. S4 for human data). The number of trials collected for the “0” and “π” conditions for each song are shown in the top right of each subpanel, respectively. An asterisk denotes conditions where the tapping distributions are significantly different between the original and pi-shifted songs (p<0.05, Watson’s U^2^ test, sample N shown in each subpanel, Bonferroni corrected for multiple comparisons), indicating that the monkey synchronized to some feature in the music when rhythmic structure was intact. **g)** Spectrograms of the temporally scrambled versions of each song in panel E used in Experiment 2. **h)** Same as panel F, but for *Experiment 2: Scrambled Music* (see Supplementary Fig. S4 for human data). Overlapping distributions suggest that the monkey produced taps based on the Go signal when temporal structure was uninformative.

To further test whether these results could have been obtained by chance, *Experiment 2: Scrambled Music* was performed whereby scrambled versions of the same three songs were presented to the monkeys (who were tasked with producing the same target intervals as in Experiment 1). Each song was scrambled by cutting it into 30 ms segments and scrambling their order using the Sound Quilting Toolbox [34] (Fig. 1g; see *Methods* and Supplementary Audio A3-A6.). These scrambled variants have the same dominant frequencies as the intact music (see Supplementary Figure S5) but perceptually lack any clear sense of a steady beat, presumably forcing the monkeys to rely on associating the sound during the Perception Phase with a target interval and producing that interval following the Go signal. As shown in Figure 1h, the monkeys can still successfully perform correct trials by producing an entirely endogenously generated beat. However, with the exception of one song for each animal, the tapping phase distributions are equivalent between the original “0” and pi-shifted “π” versions of these scrambled stimuli (p>0.05, Watson’s U^2^ test, N = 183-421 trials, Bonferroni-corrected for multiple comparisons). This indicates that while monkeys are perfectly capable of ignoring the auditory stimulus to solve the task, they choose to synchronize to some feature in the stimulus when the temporal structure is informative, as in the case of intact music (Fig. 1f).

### 2. Monkeys synchronize to beat, even when not required to

The observation that monkeys choose, seemingly spontaneously, to use features in the music to orient their tapping could be explained away as a strategy that is beneficial when tasked with producing a specific target interval. To test whether monkeys show a truly spontaneous tendency to synchronize to music (given a context of prior reward-based training), *Experiment 3: Free Tapping* was performed using a different song excerpt (‘Everybody’ by the Backstreet Boys, used previously in [20]) presented at three different tempi (552, 700, and 850 ms, see Supplementary Audio SA7-SA9), selected pseudo-randomly from trial to trial. In this experiment, monkeys initiated a trial in the same way as before, but were now completely free to produce any interval of their choosing during the tapping phase, with reward contingent only on their ITIs being internally consistent within a trial (see *Methods*). A trial ended once the monkey had produced 5 successful taps with a consistent ITI.

Figure 2a shows trials that would have been considered “correct” based on the reward contingencies of the previous experiments (ITIs within +/-20% of the true song tempo). Despite there being no reward-driven advantage to synchronizing to the music, a comparison between the original “0” and pi-shifted “π” versions of these musical excerpts indicates that, again with the exception of one case, the tapping phase distributions are significantly different when the monkey is producing ITIs that match the tempo of the song (p<0.05, Watson’s U^2^ test, N=67-485 trials, Bonferroni-corrected for multiple comparisons), indicating a spontaneous tendency to synchronize to the music even in this free tapping context.

**Fig. 2.**
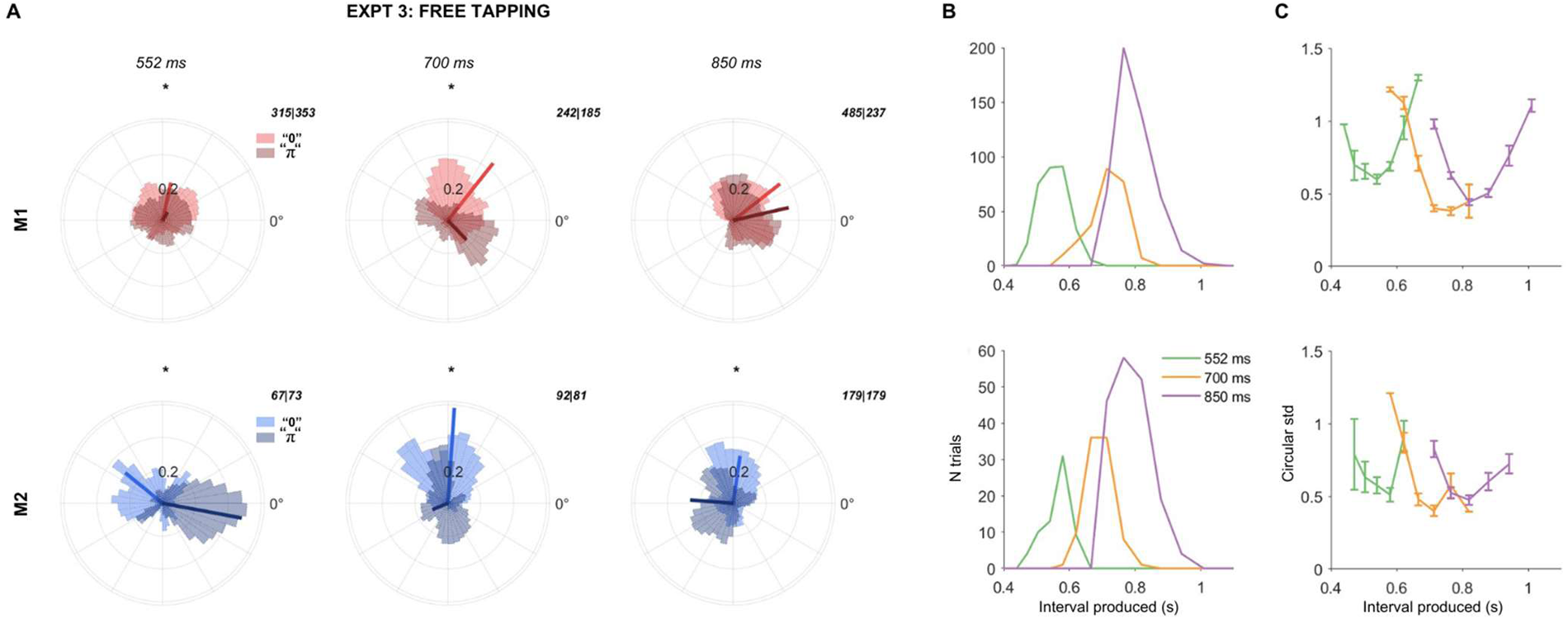
Macaques spontaneously synchronize to beat in the absence of a target interval. **a)** Tapping responses for *Experiment 3: Free Tapping* for M1 (top row) and M2 (bottom row), with song tempi of 552, 700, and 850 ms arranged from left to right. Each circular histogram shows tapping responses to the original song (in color) and pi-shifted control (same color but grayed), just for those trials where the monkey produced ITIs within +/-20% of the song tempo. 0° corresponds to the consensus tapping phase of human participants in this study for the original excerpt at the 3 respective tempi (see Supplementary Fig. S4for human data). The number of trials collected for the “0” and “π” conditions are shown in the top right of each subpanel, respectively. An asterisk denotes conditions where the tapping distributions are significantly different between the “0” and “π” conditions (p<0.05, Watson’s U^2^ test, sample N shown in each subpanel, Bonferroni corrected for multiple comparisons). Different distributions between original and pi-shifted songs indicate that the monkey synchronized to beat, even when this was not necessary for reward. **b)** Histograms of interval produced for M1 (top) and M2 (bottom), for each tempo (green: 552 ms, yellow: 700 ms, red: 850 ms). Both monkeys produce the target interval on the majority of correct trials. **c)** Circular standard deviation of tap phase on trials binned by interval produced. Circular standard deviation is lowest for trials produced at the target interval, and this pattern is consistent between M1 (top) and M2 (bottom). Colors are the same as in panel B.

Figure 2b provides insight into the intervals produced during all rewarded trials, including those where the monkey was not tapping in tempo. Here, it is clear that the peak in the distribution of ITIs is roughly at the “target” interval, even though the target interval was <u>not</u> reinforced in this experiment. This spontaneous tendency to tap at the appropriate tempo is highly consistent across both monkeys. A further striking finding is shown in Figure 2c, where each line plots the circular standard deviation of trials binned by the interval produced. Here, it is very clear that while the monkey can and often does ignore the stimulus to endogenously produce an interval of his choosing to receive reward, the tapping phases produced are maximally concentrated and consistent from trial to trial only when the monkey is tapping with ITIs that match the tempo of the song.

### 3. Comparison with humans: similar produced intervals, but differences in tapping phase

To enable direct comparison between species, 18 human volunteers performed the same experiments using the same setup as used for the monkeys (see *Methods*). Human tapping results for Experiments 1-3 are displayed in Supplementary Figure S4. Figure 3 summarizes mean and within-trial variability of tapping period (top row) and phase (bottom row) for the three original musical excerpts from Experiment 1. Individual human subjects are shown by gray dots, while M1 and M2’s respective values are shown in red and blue, respectively.

**Fig. 3.**
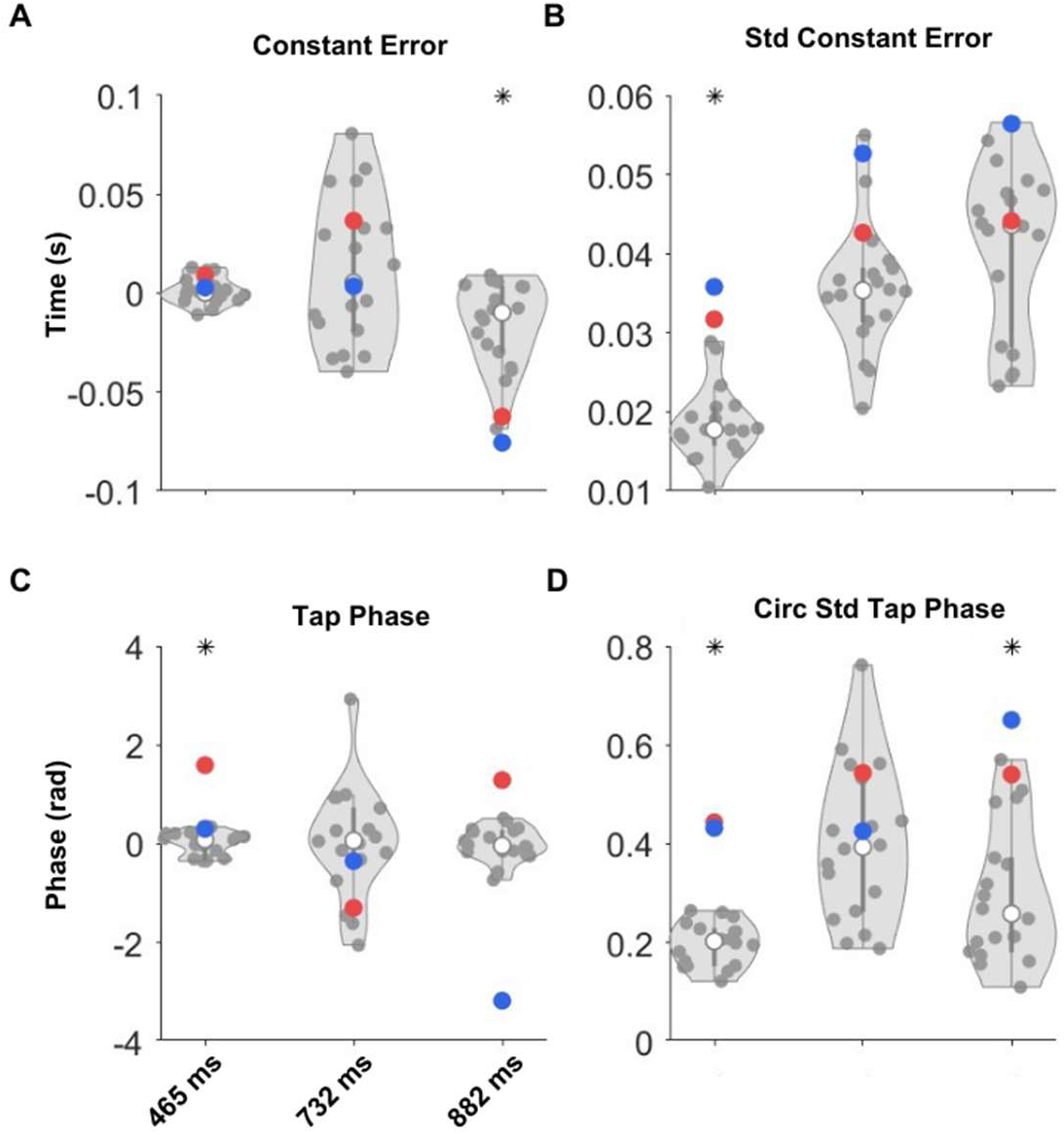
Macaques and humans produce similar beat intervals, but show differences in beat phase. Each dot represents an individual human subject (gray) or macaque (M1: red, M2: blue) based on tapping collected in *Experiment 1: Music*. **a)** Constant error, or the difference between interval produced and target interval. Positive values indicate overestimation of the beat interval. **b)** Mean standard deviation of the intervals produced within a trial for each subject. **c)** Pairwise distance between the mean interval produced between any two human subjects (gray), compared to the distance between the mean interval produced between M1 and M2 (blue). **d)** Circular mean tapping phase by subject, adjusted such that 0 represents the circular mean across human subjects. **e)** Mean of within-trial circular standard deviation of tapping phases. **f)** Pairwise distance between the circular mean tapping phase produced between any two human subjects (gray), compared to the distance between the circular mean tapping phase between M1 and M2 (blue).

Starting with tapping period, Fig. 3a indicates that both monkeys on average produced intervals that were within the range of intervals produced by humans, with the exception of the slowest song where monkeys systematically underestimated the duration of the interval (p<0.05, Watson Wilcoxon U Test, N1/N2 = 18/2 humans/monkeys). We believe this is likely due to their anxiety to receive reward [35] rather than an inability to accurately track the interval. On the other hand, monkeys produced ITIs that were more variable compared to human subjects for the fastest song (Fig. 3b), potentially indicating difficulty synchronizing to this fast tempo. However, it is worth noting the consistent trend between monkeys and humans where ITI variability increases as the interval being tapped becomes longer, consistent with the scalar property [36]. Together, these observations suggest that overall, macaques and humans show largely similar abilities to produce a target interval for each of these musical excerpts.

However, some differences emerge when examining the phase of tapping produced by monkeys versus humans. Fig. 3c shows the phases produced, relative to the “consensus” human tapping phase at 0. First, it is worth noting the much higher spread of phases produced for the intermediate tempo song even across human listeners (see also Supplementary Fig. S4). Across songs, the subjective beat phase chosen by the monkeys does not always align with that chosen by human subjects, nor does it show a systematic phase shift with respect to humans, indicating that the feature being tapped by monkeys may not correspond to the subjective human consensus beat. This is also consistent with the results in Fig. 1f where it is clear that a pi-shift in the stimulus does not produce an equivalent pi-shift in tapping responses as it (approximately) does in humans, reflecting a larger flexibility in beat interpretation. Another important difference is that there is higher variability in the phase of monkey tapping (Fig. 3d). Interestingly, monkeys and humans show similar phase variability for the more ambiguous Song 2 (see Supplementary Fig. S4 to compare the consistency of human tapping responses in Song 2 to Songs 1 and 3). This suggests that perhaps there may be a limit either to how clearly monkeys perceive or how precisely they can synchronize to musical beat, with this limit being comparable to humans tapping to a song with a less salient beat.

## Discussion

Beat perception and synchronization to music is the extraction of and synchronization to a subjective, isochronous (in the case of the musical excerpts used here) pulse from a complex rhythmic sound stream. Our data demonstrate that the macaque is capable of beat perception and synchronization. Beyond having the clear ability to do so, our macaques spontaneously chose to synchronize to music despite the possibility of obtaining reward by ignoring the sound (Experiment 2), or by producing any interval of their own choosing (Experiment 3). To our knowledge, the spontaneous coordination of auditory perception, prediction, and anticipatory action has not been previously observed in macaques. In this context, we interpret our work as uncovering a latent ability to perceive and synchronize to musical beat that expresses itself only when the component processes of auditory pattern detection, prediction, and audiomotor error correction are connected through association with reward – an idea which we frame as the “Four Components” (4Cs) hypothesis.

It is important to note that while macaques share an ability to perceive and synchronize to musical beat, we do not claim that their subjective experience of beat reflects that of humans. Unlike our human volunteers who could perform the task without training, our monkeys required extensive training and still found the task effortful, as evidenced by behaviors that reflect frustration, including immediate disengagement with the task once they consumed their needed water. Synchronization to music is therefore unlikely to be *intrinsically* rewarding to macaques. This is in contrast to humans, who willingly spend thousands of hours training in audiomotor synchronization tasks such as learning a musical instrument, a dance choreography, or even learning to speak, perhaps because of a deep and intrinsic link between the audiomotor and reward system [37–39]. On the other hand, it is likely that extensive reward-based training could confer some degree of intrinsic reward, perhaps through Pavlovian conditioning [40], which could explain why our macaques still preferred to synchronize in order to obtain reward even when other (simpler) behavioral strategies were available. We make the strong prediction that humans differ from macaques in their functional connectivity between auditory, motor, and reward regions, with structures associated with reward potentially playing a mediating role between auditory and motor processes. Future work in macaques could explore the extent to which music naturally engages motor and reward circuits as it does in humans [41], how this might change after training on a synchronization task, and whether any change is specific to the trained songs or generalizes to music that has not been previously experienced.

The quantitative differences we observed in the behavior produced by macaques and humans is also informative. Compared to humans, macaques showed a largely similar ability to tap at the 3 different target tempi, with variability increasing with longer intervals consistent with the scalar property [36], but showed larger flexibility and variability in the phase of their tapping compared to humans. This suggests that the motor system of macaques may have a similar rhythm production ability to that of humans for limb movements, but that there may be key differences in the auditory-motor loop that drive differences in the phase of tapping. This is further evidenced by previous work showing that monkeys show higher phase variability in their tapping even when presented with explicit beats from auditory metronomes [26, 28] and is consistent with the gradual audiomotor evolution hypothesis [42] that posits that the auditory-motor connections required for musical beat perception evolved gradually over primate evolution [43]. However, it is worth noting the extent of inter-individual differences even between human subjects, particularly for the more difficult medium tempo song (see Fig. 3), which has been previously reported to have the lowest tap consensus (0.81, compared to 0.88 and 0.85) of the three excerpts used here (see Supplementary Figure S6 in [33] where Songs 1, 2, and 3 correspond to Excerpts 1, 2, and 5, respectively). It appears that when humans find the beat ambiguous, the tapping for some subjects becomes as variable as that of macaques (see Supplementary Figure S4). Beyond reflecting the need for a more nuanced account of human audiomotor synchronization ability [44, 45], our findings set the stage for deeper exploration of the neural dynamics that subserve musical beat perception, guided by the similarities and differences observed in between individuals and species during synchronization to music.

A key question that will need to be addressed is why synchronization behavior differs between individuals and species. Modeling approaches may provide clues that help to pinpoint functionally where these differences are. A Bayesian framework applied to sensorimotor synchronization [46, 47] comprises 4 stages: measurement, estimation, production, and feedback, with the fitted parameters reflecting details about the underlying sensorimotor process responsible for producing the observed synchronization data. However, these models have yet to be applied to comparisons between human and macaque synchronization behavior. We believe it is likely that the monkey’s ability to estimate a Bayesian prior and produce an accurately timed movement are likely reliable, given their human-level performance during visuomotor synchronization [28]. However, their higher flexibility in tapping phase could be attributed to differences in measurement of beat, since often the same monkey produced different interpretations of beat on different trials (see Fig. 1). The higher variability in tapping phase could therefore be due to differences in either the computation of feedback (due to a less precise measurement), or in the application of accurately computed feedback (due to structural or functional limitations in audio-motor circuits). A direct comparison of fitted model parameters between human and macaque tapping to explicit metronome beats could disentangle these possibilities. From a dynamical systems perspective, beat perception could also be considered an attractor state between stimulus and tapping [48–50], where we would predict that this attractor state is stronger in humans than in macaques, affording more flexibility and variability in perception and production in the latter. Further work is required to make model-based predictions, which when tested experimentally could shed light both on inter-species and inter-individual differences in beat perception and synchronization ability.

To contextualize our finding that macaques perceive and synchronize to musical beat and to provide a wider framework for future studies, we propose the Four Components hypothesis. The 4Cs hypothesis states that animals capable of the component elements of musical beat perception may express some capacity for musical beat perception and synchronization, if doing so is rewarding. The requisite elements we propose are largely general-purpose abilities that have been reported across a wide range of species. These include an auditory component capable of abstracting salient temporal patterns out of a complex acoustic signal [33, 51–56], a predictive component capable of projecting a pattern forward to predict the timing of future stimulus input [57–62], an auditory-motor connection capable of using sensory information and feedback to time motor actions in anticipation of future sound events [63–66], and the coordination of these through and association with reward that is either intrinsic or conditioned [37–39]. This hypothesis assumes of course that ecological constraints are considered in the task design [67], for example, by choosing an appropriate effector that is capable of producing controlled movements at the desired rates. It is important to note that we do not claim or demonstrate a link here between synchronization to music and a natural occurring behavior in the macaque. Instead, we show that macaques can – through association with reward – be trained to coordinate a set of naturally occurring abilities to produce this behavior. The key advantage to this perspective is that it views musical beat perception and synchronization as a continuum – which can potentially be used to describe even the ability of different human individuals – and opens the door to comparative studies that uncover insights into human beat perception from what other species do or do not share with humans [68]. This is a clear departure from the view that only vocal learning species are capable of beat perception and synchronization [24]. However, a possible point of intersection in these theories is that auditory-motor coordination may be *intrinsically* rewarding in vocal learning species [69], whereas other in other species it would need to be associated with reward through training [70]. Future work could explore whether intrinsic functional connectivity between audiomotor and reward circuits is observed only in vocal learning species or whether this too may have evolved gradually [71, 72].

Overall, this study presents a key advance towards understanding the neurobiological and evolutionary origins of musical beat perception and synchronization. It establishes the macaque as a model organism for the study of perceiving and tapping to a subjective beat in real human music. At a theoretical level, it also provides evidence in favor of the 4Cs hypothesis which supposes that many other species may show some ability to sense musical beat, opening the study of musical beat perception to the larger range of high spatial and temporal resolution recording and manipulation techniques possible in animal models. Such techniques are already shedding light on the auditory and premotor cortical dynamics at play during sensory-motor synchronization to metronomes [30, 31, 73], which coincides with evidence from human studies that implicate auditory-motor [27, 74–76] and fronto-parietal networks [77] in audiomotor synchronization.

## Methods

### Subjects

All the animal care, housing, and experimental procedures were approved by Ethics in Research Committee of the Instituto de Neurobiología UNAM, protocol 090.A, and conformed to the principles outlined in the Guide for the Care and Use of Laboratory Animals (NIH, publication number 85-23, revised 1985). Monkeys were monitored daily by researchers and the animal care staff to check their health and well-being.

Two adult male macaques, 10 and 12 years old, participated in this study. Both macaques had been previously trained to synchronize their tapping with auditory metronomes (unpublished data) using the same hardware setup and training paradigm as used here. The monkeys were housed in separate cages on a 12-hour light/dark cycle. They were provided with dry food ad libitum, but the only water provided during their 5 days of training per week was through reward for correct trials during training and experimental sessions. Monkeys received between 200 and 250 ml of water and/or juice per day during training. If performance was especially poor or the monkey did not engage much with the task, additional water to complete the minimum 200 ml was provided through soaking dry food pellets in the required amount of water. Following the last session of the week, monkeys were given additional water up to 350 ml. They were also given 300 ml on their first rest day, and then 180 ml on their second rest day in preparation for the next day’s experimental session. The monkeys also received fruit after experimental sessions and on weekends.

Additionally, 18 human volunteers were recruited to participate in this study, including two of the authors (10 female, 2 left handed, age 21-44, median age 30; 7 with >4 years musical training). Human subjects completed one block, consisting of 30 trials, of each of the stimulus sets presented to the monkeys. In order to get accustomed to the setup, two metronome blocks were presented at the start of the session. Following this, the different stimulus sets were presented, with no explicit training provided on these stimuli.

### Training

Each monkey had been trained previously, using the same setup as used here, to tap to isochronous metronomes with intervals of 0.55 s, 0.85 s, and a range of intermediate tempi selected at random between 0.5-0.9 s. The three tempi were presented in randomized order with a ratio of 40/40/20% of trials. To facilitate training on the more complex music stimuli, care was taken to minimize the number of changes between the metronome task and the current task, while gradually training the monkeys to abstract from isolated metronome sounds (a single 50 ms tone at 1500 Hz) to continuous spectrotemporally complex music.

In the metronome task and in this training task, the monkeys would initiate a trial by holding a lever, which would simultaneously trigger the start of the auditory metronome and the appearance of a visual hold cue (yellow square) on the screen in front of the subject (the *Perception Phase*, see Fig 1). Here, the perception phase lasted four beats. The disappearance of this yellow square would signal the start of the *Production Phase* where the monkey would need to release the lever and begin tapping in the tapping area. For training, the monkey needed to produce a series of 6 taps, each with an asynchrony within +/-0.23 s of sound onsets, in order to receive reward at the end of the trial. If a tap asynchrony exceeded this window, the trial would abort immediately, which was signaled by the cessation of further sounds. In general, a minimum delay of 5 seconds would pass before the next trial could be initiated after either a successful or error trial, but this value was adjusted manually up or down by the experimenter depending on the motivation level of the monkey.

At the start of training for this study, the metronome task was applied but replacing the metronome tone with a 50 ms tone cloud (see *Stimuli*). As the monkey adjusted to synchronizing to the new, spectrotemporally complex sound, the duration of the tone cloud was lengthened by another 50 ms, such that it occupied a larger portion of the inter-stimulus interval. The tone cloud was further lengthened, each time by 50 ms, over a variable number of sessions until the tone clouds were 450 ms in duration. At this stage, the 450 ms tone cloud occupied nearly the entirety of the shortest possible inter-stimulus interval of 500 ms.

Once the monkey reliably performed the metronome task with 450 ms tone clouds, continuous tone clouds that repeated seamlessly at three fixed tempi were presented (550, 700, and 850 ms, with a ratio of 40/20/40%; see *Stimuli*). The repetition of these random stimuli induces a sense of subjective beat that can be tapped (see [78] for human tapping to seamlessly repeating white noise). Importantly, since there are no longer clear onsets, and because synchronization to these continuous stimuli depends on a subjective judgment of beat, reward here was made contingent on the produced inter-tap intervals (IP) being within +/-30% of the target interval. Initially, the monkeys were required to produce fewer taps, beginning with 3 taps. As the monkey improved in this task, the number of taps was increased incrementally to 6, while the tolerance window for produced intervals was gradually narrowed to a final +/-20% of the target interval. In a given session, only three unique stimuli were presented, one at each tempo. However, over training, different sets of tone clouds were also presented to ensure that monkeys were able to generalize the task to different exemplars of the same class of stimuli.

Once the monkey reached a reliable level of performance on the continuous tone clouds, the monkey was considered trained. Thereafter, the first sessions in which the final experimental stimuli were presented were conducted with the final parameters, with the exception of the number of taps that needed to be produced. Here, like in the training sessions, monkeys began by needing to produce 3, then 4, and then finally 5 or 6 taps. The first 5 taps of all correct trials that had a minimum of 5 produced taps are analyzed here.

Due to the effortful nature of the task for the monkeys, the criterion to determine a reliable level of performance was not fixed but was determined by the judgment of the experimenter based on the need to balance the motivation of the monkey with advancement in training. Signs of reliability included an overall percentage of correct trials >40%, with intermittent spells of accuracy >60% within a session, over at least two consecutive sessions. If in a session the monkey’s performance was very poor, a step back was taken in the training procedure, for example by requiring one fewer tap, until performance stabilized again.

A session was continued until the monkey lost motivation to do the task. Usually, this occurred after the monkey had consumed 200-250 mL of water. If the monkey disengaged before he had consumed sufficient water, the juice was provided as reward instead of water, which often motivated the monkey to continue working. A monkey produced between 150 and 300 trials in a session, depending on his motivation and level of comfort with the task and stimuli.

### Stimuli

#### General task

To avoid the possibility that the monkeys could employ a strategy seeking sound content at 150 Hz (corresponding to the frequency of the metronome tone they had been previously trained on), and to match the frequency range of the tone clouds employed during training, all musical excerpts were bandpass filtered between 200 and 6400 Hz with a stopband attenuation of 120 dB.

Three different stimuli, each with a different tempo, were presented in a pseudo-randomly selected manner within a given session. The fast, medium, and slow stimuli were presented on 40/20/40% of trials, respectively. The stimulus for each trial was selected randomly, but out of mini-batches of 5 trials containing 2/1/2 trials of each stimulus. This was done so that monkeys were obliged to produce correct trials for all stimuli. This meant that occasionally, the same trial could be repeated until the monkey synchronizes correctly, advancing to the next mini-batch containing all stimuli.

#### Training: Tone clouds

Tone clouds are a class of random sounds parametrized by two parameters: tones per second and tones per octave [79]. The repetition of even arbitrary, random sounds such as white noise is easily detected by humans [80, 81] and also generates a pulse that can be tapped by listeners [78, 82]. Tone clouds can be sparse or dense, depending on the number of tones/second/octave, and at high densities are indistinguishable from white noise [79]. Here, relatively sparse tone clouds were used.

Based on generating a “grid” with the specified tones/sec and tones/oct parameters (see Fig. 1 in [79]), a tone onset is placed inside the confines of each cell with a random frequency and a random latency. The tone clouds used here had a spectral density of 1 tone/oct (over the 6 octaves between 200 Hz and 6400 Hz) and a temporal density of 2 tones/s. Due to the target intervals from the metronome task of 550, 700, and 850 ms, a temporal grid of 25 ms was chosen as the least common multiple of these values. To approximate the desired temporal density of 2 tones per second, each “cell” along the temporal grid had a probability of 1/20 of containing a tone, with tone frequency and latency chosen randomly within the cell. Tones were 50 ms in duration and could overlap with other tones from other cells in the grid.

#### Experiment 1: Musical excerpts and pi manipulation

The first 10 seconds of Excerpts 1, 2, and 5 from the MIREX 2006 dataset [83] and used previously in [33] were used here (**Song 1:** *‘You’re My First, My Last, My Everything’* by Barry White; **Song 2:** *‘A New England’* by Billy Bragg; **Song 3:** *‘Passe & Medio Den Iersten Gaillar’* by Josquin Des Prez, respectively). These songs were chosen based on their tempi being close to the metronome tempi the monkeys had been trained to tap to previously (550, 700, 850 ms). They were also chosen based on having a relatively high tapping consensus across humans (0.88, 0.81, and 0.85; see Excerpts 1, 2, and 5 in Supplementary Figure S6 from [33]) in an attempt to make the task as easy as possible for the monkeys. The Perception Phase for musical excerpts included 1 s plus 4 beat intervals, for consistency with the training with tone clouds.

To generate the pi-shifted stimuli, the start of each musical excerpt was truncated by a duration corresponding to half of one beat interval (465/2, 732/2, and 850/2 ms; see Fig. 1c). Each pi-shifted stimulus therefore started with features that happened half of a beat interval *later* in the original stimuli. However, the visual go-cue still occurred 1s plus 4 beat intervals into the pi-shifted stimuli. Thus, subjects were required to begin tapping at a pi-shifted location with respect to the features in each song.

#### Experiment 2: Scrambled musical excerpts

The three bandpassed original excerpts from Experiment 1 were subjected to the quilting procedure from. Briefly, the Sound Quilting Toolbox cuts a sound signal into segments of a given duration (here, 30 ms segments were specified), shuffles them, and reconstructs the signal without any audible artifacts. The resulting scrambled versions of each musical excerpt were then pi-shifted by truncating from their respective starts, as described earlier.

#### Experiment 3: Same song 3 tempi

In this experiment, “Everybody” by the Backstreet Boys was used. The original tempo of the song was 552 ms. Additional tempi of 700 and 850 ms were generated by slowing down the original bandpassed excerpt without changing the pitch using the Matlab function *stretchAudio*.

### Analysis

#### Data processing

All trials containing at least 5 correct taps were analyzed here. For Experiment 1 and 2, taps were considered correct if the inter-tap intervals between taps 2-5 were within +/-20% of the target interval for that trial. For Experiment 3 where the monkey was not constrained by a target interval, a trial was considered correct if taps 3-5 had inter-tap intervals that were within +/-20% of the monkey’s own inter-tap interval between taps 2 and 3. No further processing or exclusion of data was done.

The concentration of taps within a beat interval was determined by expressing tap times in radians. Given the subjective nature of beat, 0° was set to be the circular mean based on human tapping for each stimulus. This was done by computing the circular mean for humans, and subtracting this value from all tap times across humans and monkeys. Thus, the tapping phases produced by each subject are interpretable as being relative to the consensus beat tapped by our pool of human subjects. For visualization, circular histograms were smoothed with a kernel width of pi/20 using the Matlab function *circ_ksdensity* [84].

#### Statistical analyses

The Watson’s U^2^ test for circular distributions was used to determine whether the tapping phases for the original and pi-shifted stimuli differed significantly (Matlab function *watsons_U2* [85]). This nonparametric test is appropriate due to the multimodal distributions produced by the animals. A p-value cutoff of 0.05 was selected, with Bonferroni correction for multiple comparisons (2 monkeys x 3 stimuli).

Statistical comparisons between monkey and human data were done using the Mann Whitney Wilcoxon Test (Matlab function *mwwtest* [86]), a nonparametric test for effects between two groups of unequal and possibly small sizes. A cutoff of p<0.05 was selected, without correction for multiple comparisons due to the number of monkey samples (N=2) being at the lower limit of the statistical tables for the U statistic.

## Supporting information

Supplementary Figures and Media

## Acknowledgements

We are grateful for the valuable comments that Victor de Lafuente and Nori Jacoby provided to our manuscript. We also thank Maria Antonieta Carbajo, Juan Ortíz, and Raul Paulín, for their technical assistance. Supported by CONACYT: A1-S-8430, PAPIIT: IG200424, and UNAM-DGAPA-PASPA.

