## Supplementary Figures and Media for "Monkeys have rhythm"

### Supplementary Material

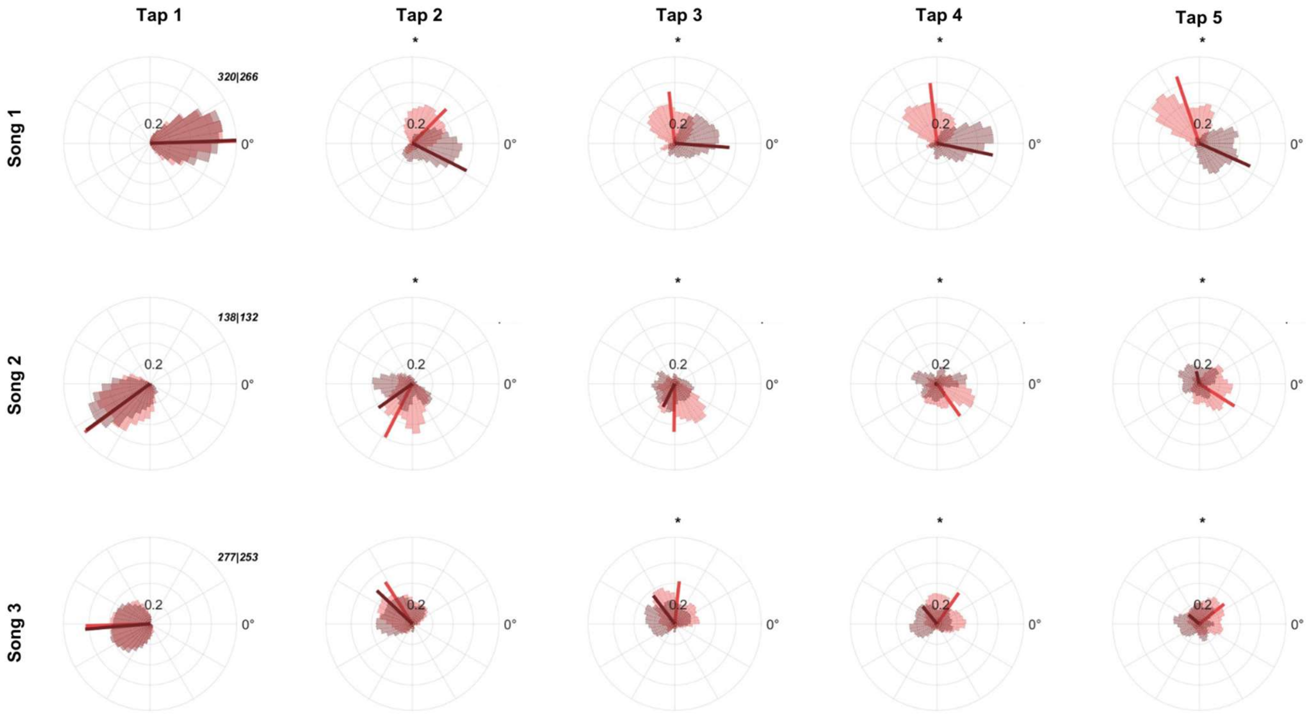


**Fig. S1. Tapping between original and pi-shifted songs diverges by 2nd or 3rd tap for monkey M1.** Rows are Songs 1-3 (top to bottom, respectively), and columns are Taps #1-5, arranged from left to right. An asterisk denotes conditions where the tapping distributions are significantly different between the original and pi-shifted songs for the given tap in the sequence ($p<0.05$, Watson's $U^2$ test, sample N shown in the Tap 1 subpanel, Bonferroni corrected for multiple comparisons).

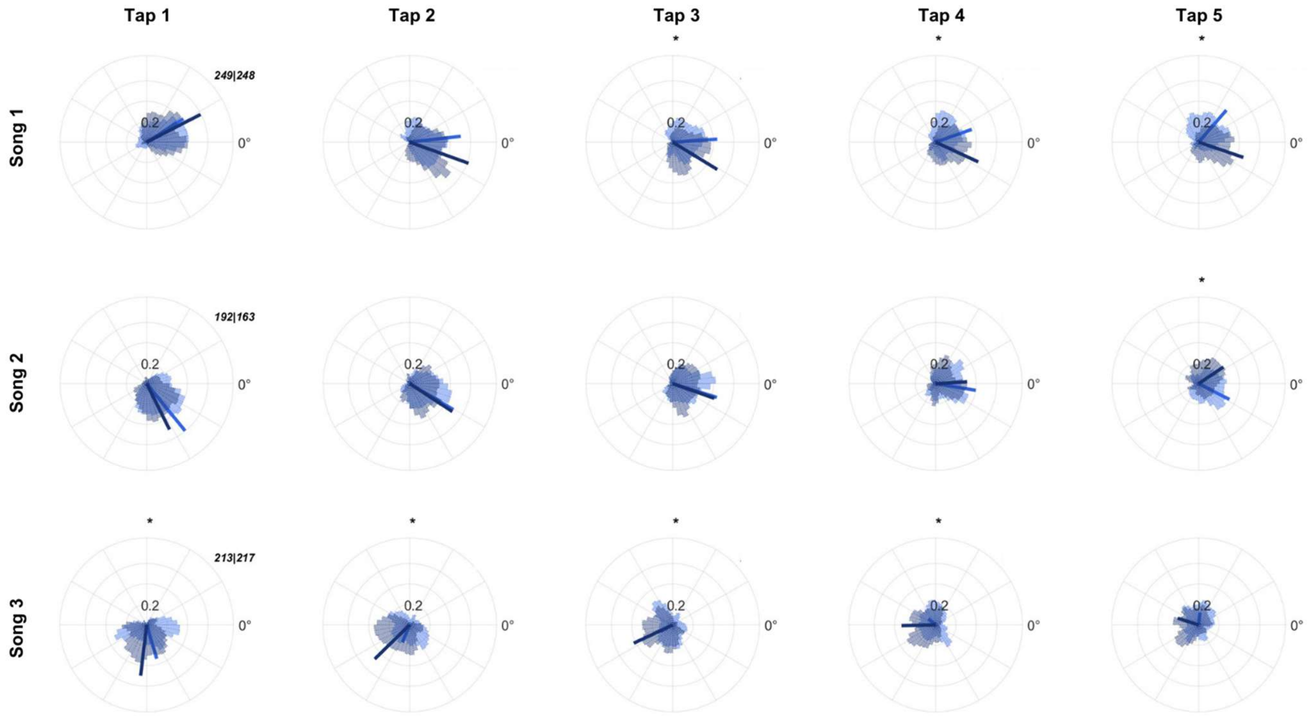


**Fig. S2. Tapping between original and pi-shifted songs diverges for Song 1 and Song 3 for monkey M2.** Rows are Songs 1-3 (top to bottom, respectively), and columns are Taps #1-5, arranged from left to right. An asterisk denotes conditions where the tapping distributions are significantly different between the original and pi-shifted songs for the given tap in the sequence ($p<0.05$, Watson's $U^2$ test, sample N shown in the Tap 1 subpanel, Bonferroni corrected for multiple comparisons).

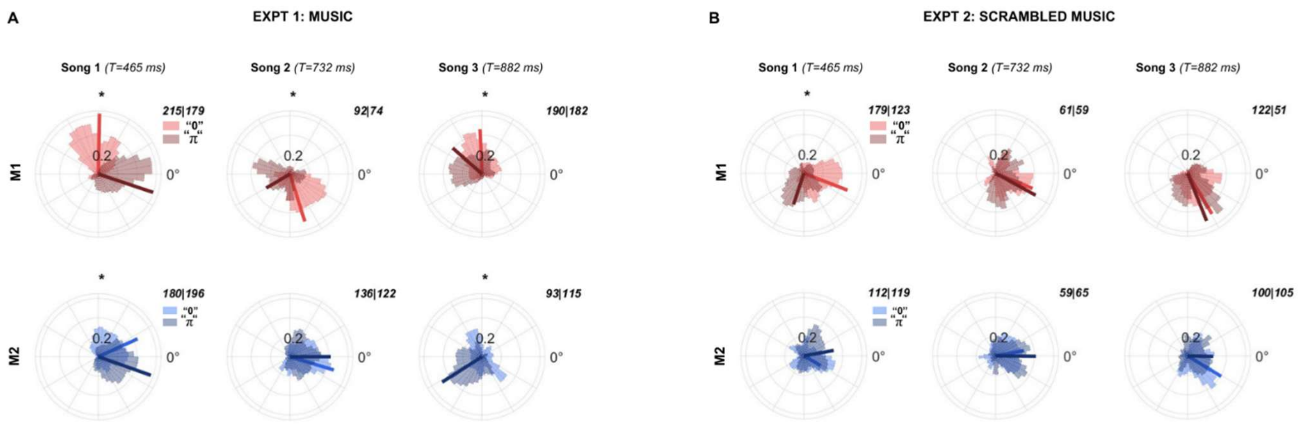


**Fig. S3. Macaque tapping during higher quality trials in Experiments 1-2.** Data are for all trials where each produced interval is within +/- 15% of the target interval (as opposed to 20% in the main figures). **A)** Data for Experiment 1 using music excerpts, compare with main Fig. 1F. **B)** Data for Experiment 2 with scrambled music excerpts, compare with main Fig. 1H. Note that the difference between "0" and "π" reported in Fig. 1H for M2 in Song 3 is insignificant here ($p>0.05$, Watson's $U^2$ test, Bonferroni corrected for multiple comparisons).

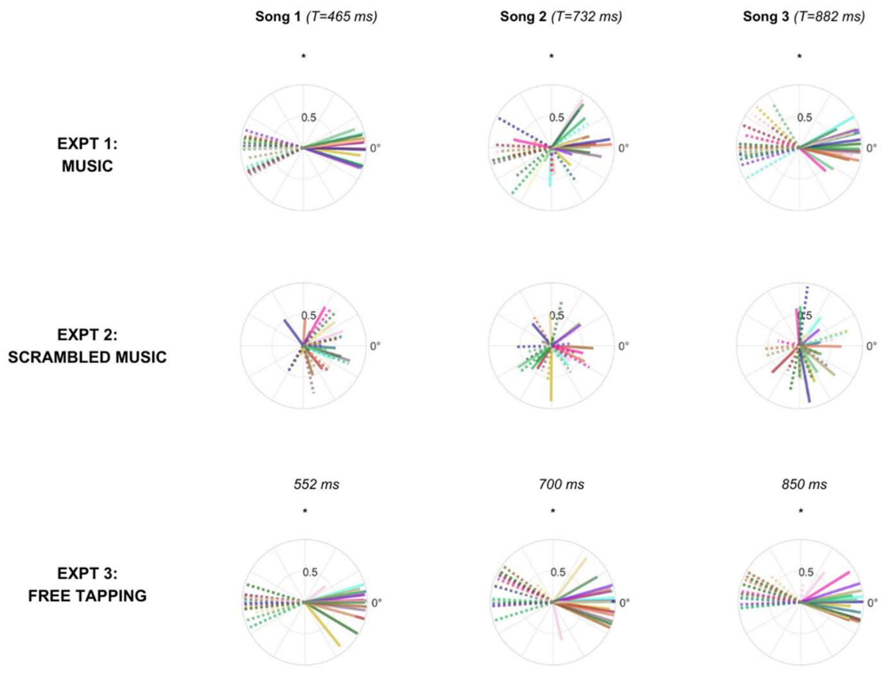


**Fig. S4. Summary of human tapping for Experiments #1-3.** Rows are Experiments 1-3, and columns are Songs #1-3, arranged from left to right. Each subject is represented a their resultant vector in a different color (N=18), with solid lines showing a subject's tapping phase interpretation for the original song "0", and the dotted line in the respective color representing that subject's pi-shifted control data "π". An asterisk denotes conditions where the tapping distributions are significantly different between the original and pi-shifted songs for the given tap in the sequence ($p<0.05$, Watson's $U^2$ test, N=18 subjects, Bonferroni corrected for multiple comparisons). Note that for Experiment 1, the difference between the "0" and "π" conditions is significant for all three songs, but is not always a difference of exactly π. Human tapping to the medium tempo song is particularly variable, as evidenced by less consistency in tapping phase between individuals, and shorter resultant vectors that indicate higher phase variability within a subject. In Experiment 2, no significant differences between "0" and "π" are observed, indicating that listeners are not synchronizing their taps to any feature in the scrambled music. In Experiment 3, the differences between "0" and "π" are significant for all three song tempi.

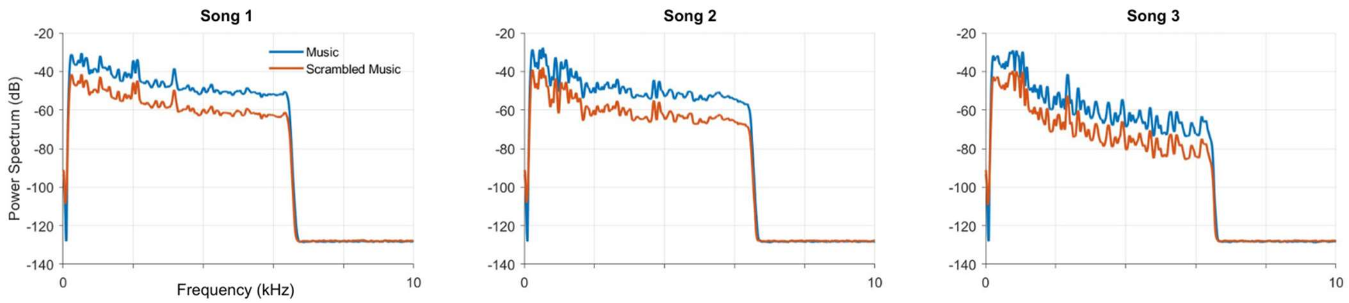

**Fig. S5. Power spectra of musical excerpts and their scrambled variants (Experiments 1 & 2).** Dominant frequencies in each song are highly similar despite the scrambling procedure.

#### Audio files

The files used in the experiment are listed below and are accessible from this link:
https://www.dropbox.com/scl/fo/n88q0u5tt59dfshicbrnm/h?rlkey=ia6s2ih82tk0ydx8bksa17mq2&dl=0

**Audio A1. *Experiment 1: Music*, Song 1, tempo 465 ms:** *'You're My First, My Last, My Everything'* by Barry White
**Audio A2. *Experiment 1: Music*, Song 2, tempo 732 ms:** *'A New England'* by Billy Bragg
**Audio A3. *Experiment 1: Music*, Song 3, tempo 882 ms:** *'Passe & Medio Den Iersten Gaillar'* by Josquin Des Prez

**Audio A4. *Experiment 2: Scrambled Music*,** Scrambled version of Song 1 from Experiment 1
**Audio A5. *Experiment 2: Scrambled Music*,** Scrambled version of Song 2 from Experiment 1
**Audio A6. *Experiment 2: Scrambled Music*,** Scrambled version of Song 3 from Experiment 1

**Audio A7. *Experiment 3: Free Tapping*, Tempo 1: 552 ms,** *'Everybody'* by Backstreet Boys
**Audio A8. *Experiment 3: Free Tapping*, Tempo 2: 700 ms,** *'Everybody'* by Backstreet Boys
**Audio A9. *Experiment 3: Free Tapping*, Tempo 3: 850 ms,** *'Everybody'* by Backstreet Boys

#### Video files

**Video V1.** Monkey M1 performing an example correct trial for Songs #1-3 in *Experiment 1: Music*.
**Video V2.** Monkey M2 performing an example correct trial for Songs #1-3.
**Video V3.** Monkey M2 performing two different interpretations of beat for Song #2.
